# The histone chaperone FACT facilitates heterochromatin spreading through regulation of histone turnover and H3K9 methylation states

**DOI:** 10.1101/2021.06.30.450523

**Authors:** Magdalena Murawska, R. A. Greenstein, Tamas Schauer, Karl C.F. Olsen, Henry Ng, Andreas G. Ladurner, Bassem Al-Sady, Sigurd Braun

**Affiliations:** Biomedical Center, Physiological Chemistry, Ludwig-Maximilians-University of Munich, 82152 Planegg-Martinsried, Germany; Department of Microbiology and Immunology and the George Williams Hooper Foundation, University of California San Francisco, San Francisco CA 94143, United States; TETRAD graduate program, University of California San Francisco, San Francisco CA 94143, United States; Biomedical Center, Bioinformatics Unit, Ludwig-Maximilians-University of Munich, 82152 Planegg-Martinsried, Germany; International Max Planck Research School for Molecular and Cellular Life Sciences, Planegg-Martinsried, Germany

**Author notes:** Correspondence (M.M.), (B. A-S.), (S.B.).

**Keywords:** FACT, histone chaperone, heterochromatin spreading, Epe1, histone turnover

## Abstract

Heterochromatin formation requires three distinct steps: nucleation, self-propagation (spreading) along the chromosome, and faithful maintenance after each replication cycle. Impeding any of those steps induces heterochromatin defects and improper gene expression. The essential histone chaperone FACT has been implicated in heterochromatin silencing, however, the mechanisms by which FACT engages in this process remain opaque. Here, we pin-pointed its function to the heterochromatin spreading process. FACT impairment reduces nucleation-distal H3K9me3 and HP1/Swi6 accumulation at subtelomeres and derepresses genes in the vicinity of heterochromatin boundaries. FACT promotes spreading by repressing heterochromatic histone turnover, which is crucial for the H3K9me2 to me3 transition that enables spreading. FACT mutant spreading defects are suppressed by removal of the H3K9 methylation antagonist Epe1 via nucleosome stabilization. Together, our study identifies FACT as a histone chaperone that specifically promotes heterochromatin spreading and lends support to the model that regulated histone turnover controls the propagation of epigenetic marks.

---

The eukaryotic genome is partitioned into transcriptionally active euchromatin and transcriptionally silent heterochromatin. Heterochromatin is instrumental for genome protection, proper chromosome segregation and maintenance of cell-type specific gene expression patterns (Becker et al., 2016; Janssen et al., 2018; Penagos-Puig and Furlan-Magaril, 2020).

Fission yeast (*Schizosaccharomyces pombe*) is a powerful model organism to study heterochromatin formation and inheritance (Allshire and Ekwall, 2015; Goto and Nakayama, 2012; Mizuguchi et al., 2015). Heterochromatin is present in this organism at distinct chromosomal regions, including repetitive sequences at pericentromeres and subtelomeres, the silent mating-type locus and facultative heterochromatin islands (Allshire and Ekwall, 2015; Wang et al., 2014; Zofall et al., 2012). Heterochromatin is characterized by the presence of hypoacetylated and H3K9-methylated nucleosomes, which are bound by the HP1 family chromodomain proteins Swi6 and Chp2. This heterochromatin platform recruits other accessory factors to safeguard transcriptional and post-transcriptional gene silencing (Ekwall et al., 1995; Reyes-Turcu and Grewal, 2012; Yamada et al., 2005).

Heterochromatin assembly initiates at nucleation sites where the RNAi machinery, cis-DNA sequences, or the shelterin complex recruit the sole histone 3 (H3) lysine 9 (K9) methyltransferase, Clr4 (Martienssen and Moazed, 2015; van Emden et al., 2019; Wang et al., 2016; Wang and Moazed, 2017). While Clr4 modifies K9, it can also bind the K9 methylation mark through its chromodomain. This allows for a ‘write’ and ‘read’ mechanism of heterochromatin establishment promoting the self-propagation of this repressive mark as well as its re-establishment after each cell division (Chen et al., 2008; Zhang et al., 2008). A differential affinity of Clr4 and Swi6 chromodomains towards H3K9me2 and H3K9me3, respectively, is believed to be important for efficient heterochromatin spreading from nucleation sites over large distances (Al-Sady et al., 2013; Jih et al., 2017; Zhang et al., 2008). heterochromatin spreading leads to generation of heterochromatic regions, which are essential for genome stability and cell fate maintenance (Greenstein and Al-Sady, 2019). While the heterochromatin nucleation has been extensively studied, the mechanisms of heterochromatin expansion along chromosome arms remain obscure.

The histone chaperone FACT (FAcilitates Chromatin Transcription) is an essential and highly conserved protein complex with two subunits, Spt16 and Pob3/SSRP1. FACT has been extensively studied in the context of actively transcribed genes where it facilitates nucleosome dis- and re-assembly in the wake of RNA Polymerase II (Duina et al., 2007; Formosa and Winston, 2020; Lee et al., 2017; Murawska et al., 2020; Orphanides et al., 1998). In addition, FACT has been implicated in a plethora of other chromatin associated processes, such as DNA repair and replication (Herrera-Moyano et al., 2014) or higher order chromatin formation (Garcia-Luis et al., 2019; Murawska et al., 2020).

Beyond its functions in euchromatic transcription, we showed previously that FACT is also involved in pericentromeric heterochromatin silencing, acting in a pathway independent of the RNAi machinery (Lejeune et al., 2007). A recent study further showed that FACT interacts with a perinuclear protein complex that facilitates FACT loading on the mating-type heterochromatin to suppress histone turnover and promote heterochromatin maintenance (Holla et al., 2020). However, the exact function of FACT in heterochromatin formation needs to be established. Specifically, whether FACT predominantly acts during heterochromatin nucleation, spreading and/or maintenance phases remains unknown.

Here, we determined a specific function of FACT in the heterochromatin formation process. We found that transcriptional silencing is impaired throughout heterochromatin in mutants deficient in FACT. However, the heterochromatin structure is particularly affected at heterochromatin-euchromatin transitions, which display reduced heterochromatin domain expansion suggesting a heterochromatin spreading defect. By monitoring heterochromatin propagation at the single-cell level, we confirmed that FACT is critical for heterochromatin spreading. We provide further evidence that FACT limits heterochromatic histone turnover, which is critical for facilitating a productive K9 tri-methylation step by Clr4. Deletion of the Jumanji protein Epe1, but not other factors antagonizing heterochromatin, suppresses the spreading defects of FACT by stabilizing heterochromatic nucleosomes. Together, our data reveal an unexpected function of the highly conserved FACT complex in heterochromatin spreading along chromosomal arms.

## Results

### Transcriptional gene silencing is impaired at different levels at pericentromeres and subtelomeres in the FACT mutant

In order to understand the functional contribution of FACT to heterochromatin silencing, we sought to systematically analyze and compare heterochromatin structure and gene repression at the genome-wide level. For this analysis we utilized the *pob3Δ* mutant, which in contrast to other model organisms is not lethal but recapitulates many of the known FACT defects (Murawska et al., 2020). In agreement with previous studies, ChIP-seq revealed only subtle change of H3K9me2 at pericentromeres (Fig. 1A, 1B and S1D) (Holla et al., 2020; Lejeune et al., 2007). This result suggests that heterochromatin domains strictly depending on RNAi are not impaired when FACT is depleted. In contrast to pericentromeres, there was a substantial reduction of H3K9me2 at the mating-type locus, in agreement with previous study (Holla et al., 2020) (Fig. S1A), and at the subtelomeres of chromosomes I and II (Fig. 1A). Interestingly, H3K9me2 was more affected in *pob3Δ* at the distal subtelomeric heterochromatin than at the telomeric ends where heterochromatin nucleation takes place. Moreover, rDNA and several facultative heterochromatin islands had also reduced H3K9me2 levels in the *pob3Δ* strain (Fig. S1B and S1C).

**Figure 1.**
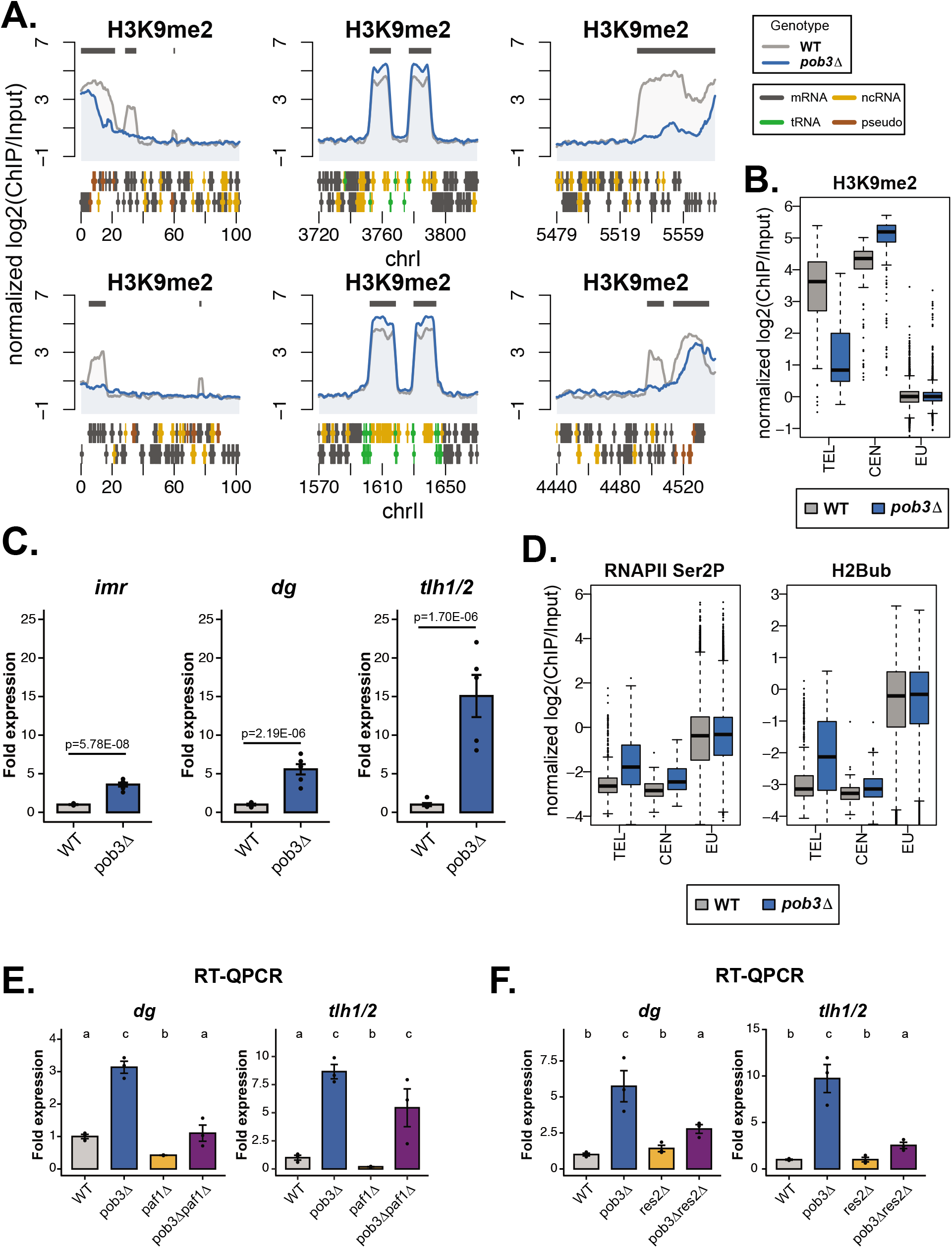
Transcriptional silencing is impaired in the FACT mutant. A. H3K9me2 ChIP-seq enrichment [normalized log2(ChIP/Input)] at subtelomeres and pericentromeres in WT and *pob3Δ* strains on chromosomes I and II. Colors: grey – WT, blue – *pob3Δ*. Both (+) and (−) DNA strands are shown with the indicated gene coding (mRNA), noncoding (ncRNA), tRNA and pseudogenes. Dark grey bars over the graphs indicate the localization of H3K9me2 in the WT strain. B. Boxplots of H3K9me2 ChIP-seq enrichment [normalized log2(ChIP/Input)] calculated in 250 bp bins in WT and *pob3Δ* strains. TEL – subtelomeres, CEN – pericentromeres, EU – rest of the genome (euchromatin). Average of 2 biological replicates is shown. C. RT-QPCR analysis. Expression of heterochromatin transcripts (pericentromeric - *imr*, *dg* and subtelomeric - *tlh1/2*) in the *pob3Δ* strain relative to WT after normalization to *act1*+. n=5-6 biological replicates; data are represented as mean +/-SEM. Statistical analysis was done on log2 transformed values. One-way ANOVA was performed and p-values were calculated with a Tukey’s post hoc test at P < 0.05. D. Boxplots of RNAPII Ser2P and H2Bub ChIP-seq enrichment [normalized log2(ChIP/Input)] calculated in 250 bp bins in the WT and *pob3Δ* strains. Average of 2 biological replicates is shown. Labeling as in Figure 1B. E. and F. RT-QPCR analysis. Expression of heterochromatin transcripts (*dg* and *tlh1/2*) in the *pob3Δ*, *paf1Δ* and *pob3Δpaf1Δ* (E.) or *pob3Δ*, *res2Δ* and *pob3Δres2Δ* (F.) relative to WT after normalization to *act1*+. n=3 biological replicates; data are represented as mean +/-SEM. Statistical analysis was done on log2 transformed values. One-way ANOVA was performed, different letters denote significant differences with a Tukey’s post hoc test at P < 0.05. See also Figure S1.

Next, we analyzed if changes in the H3K9me2 levels are accompanied by de-repression of heterochromatic transcripts in *pob3Δ*. RT-QPCR analysis showed substantial accumulation of subtelomeric transcripts but also, albeit to a lower degree, increase in pericentromeric transcripts (Fig. 1C). The absence of strong pericentromeric defects suggested a role of FACT in transcriptional (TGS) rather than post-transcriptional gene silencing (PTGS). Hence, we analyzed the abundance of the active transcription machinery at heterochromatin in *pob3Δ* using previously published ChIP-seq datasets (Murawska et al., 2020). In analogy to the H3K9me2 analysis (Fig. 1B), we divided the genome into subtelomeres (TEL), pericentromeres (CEN) and euchromatin (EU). At subtelomeres, but not at euchromatin, we found increased signals of the elongating form of RNA polymerase II (RNAPII Ser2P) and H2B ubiquitination (H2Bub), a histone mark associated with active transcription (Fig. 1D and S1D). RNAPII Ser2P and H2Bub were also slightly enriched at pericentromeres, in agreement with our heterochromatin expression analysis (Fig. 1C and 1D). Moreover, deletion of factors involved in transcription elongation (*paf1+*) or termination (*res2+*), alleviated the *pob3Δ* silencing defect at pericentromeric and subtelomeric heterochromatin (Fig. 1E and 1F). Together, our data suggest that TGS is impaired in the *pob3Δ* mutant at the genome-wide scale. In particular, sites distal to heterochromatin nucleation sites are most vulnerable to FACT depletion.

To gain a more comprehensive picture of FACT engagement with heterochromatin, we examined its genome-wide distribution by comparing the enrichment of the two FACT subunits, Pob3 and Spt16, at subtelomeres, pericentromeres, and euchromatin (Fig. S1E). This analysis showed highest levels of FACT at euchromatin, lower at subtelomeres and the lowest at pericentromeres. This result is in accordance with recent models supporting enhanced FACT recruitment via distorted nucleosomes, e.g., at transcribed regions (Liu et al., 2020). Further analysis revealed that despite the low levels at perincentromeres, FACT accumulates at the core centromeres of all *S. pombe* chromosomes, which are transcribed but largely void of H3K9me2 (Fig. S1E) (Sadeghi et al., 2014). Thus, we conclude that FACT accumulates only at relatively low levels at heterochromatic regions, which is likely sufficient to maintain heterochromatin functions.

Altogether, our genome-wide analysis of FACT silencing defects suggests that FACT’s contribution to heterochromatin maintenance is less critical at regions containing frequent nucleation sites, like pericentromeres. However, its role becomes more important at nucleation-distal regions and particularly towards heterochromatin boundaries.

### FACT controls heterochromatin spreading

Although H3K9me2 was partially reduced at subtelomeres, we found that Clr4 binding was unaltered (Fig. S2A). This result suggests that heterochromatin nucleation is not affected in the FACT mutant. The gradual reduction of H3K9me2 at subtelomeres (Fig. 1A) prompted us to investigate whether FACT plays a role instead in heterochromatin spreading. Precise analysis of heterochromatin spreading requires the ability to record heterochromatin assembly quantitatively both at nucleation and distal sites. To this end, we applied a recently developed single-cell heterochromatin spreading sensor (HSS) assay (Greenstein et al., 2018). In this system, two fluorescent reporters are integrated at different sites to report separately on heterochromatin nucleation (green) and spreading (orange). An additional red reporter in an unrelated euchromatic locus is used to filter for a cellular noise (Fig. 2A). We integrated the HSS reporter system at the silent mating-type locus, since this chromatin region has been widely used for studying heterochromatin spreading mechanisms (Greenstein et al., 2018; Holla et al., 2020; Shipkovenska et al., 2020). For our analysis, we applied a genetic background where the spreading sensor reporter was introduced in the *mat* locus with a mutationally inactivated *REIII* element, which is deficient in recruiting Clr4, HP1 and histone deacetylases via the transcription factors Atf1 and Pcr1 (Fig. 2B) (Jia et al., 2004). Hence, in this strain background, H3K9me heterochromatin nucleation and spreading is driven exclusively from the RNAi-pathway-dependent *cenH* element. To quantitatively assess heterochromatin formation, flow cytometry (FC) was performed on log-phase cells (Greenstein et al., 2018). Importantly, FC analysis showed no obvious changes in the cell cycle profiles of mutant deficient in FACT activity in contrast to the G1/M-arrested *ts cdc25* cells, which served as a control for cell cycle perturbations (Fig. S2B). For the control *clr4Δ* strain, in which H3K9 methylation is completely erased, the HSS assay showed derepression of both green and orange reporters compared to the WT strain (Fig. 2C). Conversely, in the *pob3Δ* strain, the orange reporter was also fully derepressed in the majority of cells, yet the green reporter remained silenced or mildly derepressed compared to WT. This result implies that *pob3Δ* cells have a heterochromatin spreading defect (Fig. 2C). We next examined whether the other FACT subunit Spt16 had similar effects. Unlike *pob3Δ*, *spt16* deletions are inviable. Hence, we used the temperature sensitive *spt16-1* allele containing several point mutations in the Spt16 N-terminal domain that affect FACT stability at the restrictive temperature (Choi et al., 2012). While *spt16-1* cannot grow at 32°C, it is viable at 27°C. We first assessed whether FACT activity or protein levels are affected at the non-restrictive temperature. Western blot and ChIP analysis revealed a substantial reduction of both Spt16 and Pob3 protein levels and their presence at euchromatin and heterochromatin at 27°C (Fig. S2C, S2D and S2E). We concluded that the *spt16-1* mutant is a partial loss of FACT function allele when grown at 27°C, which provides a unique opportunity to study specific function(s) of this histone chaperone.

**Figure 2.**
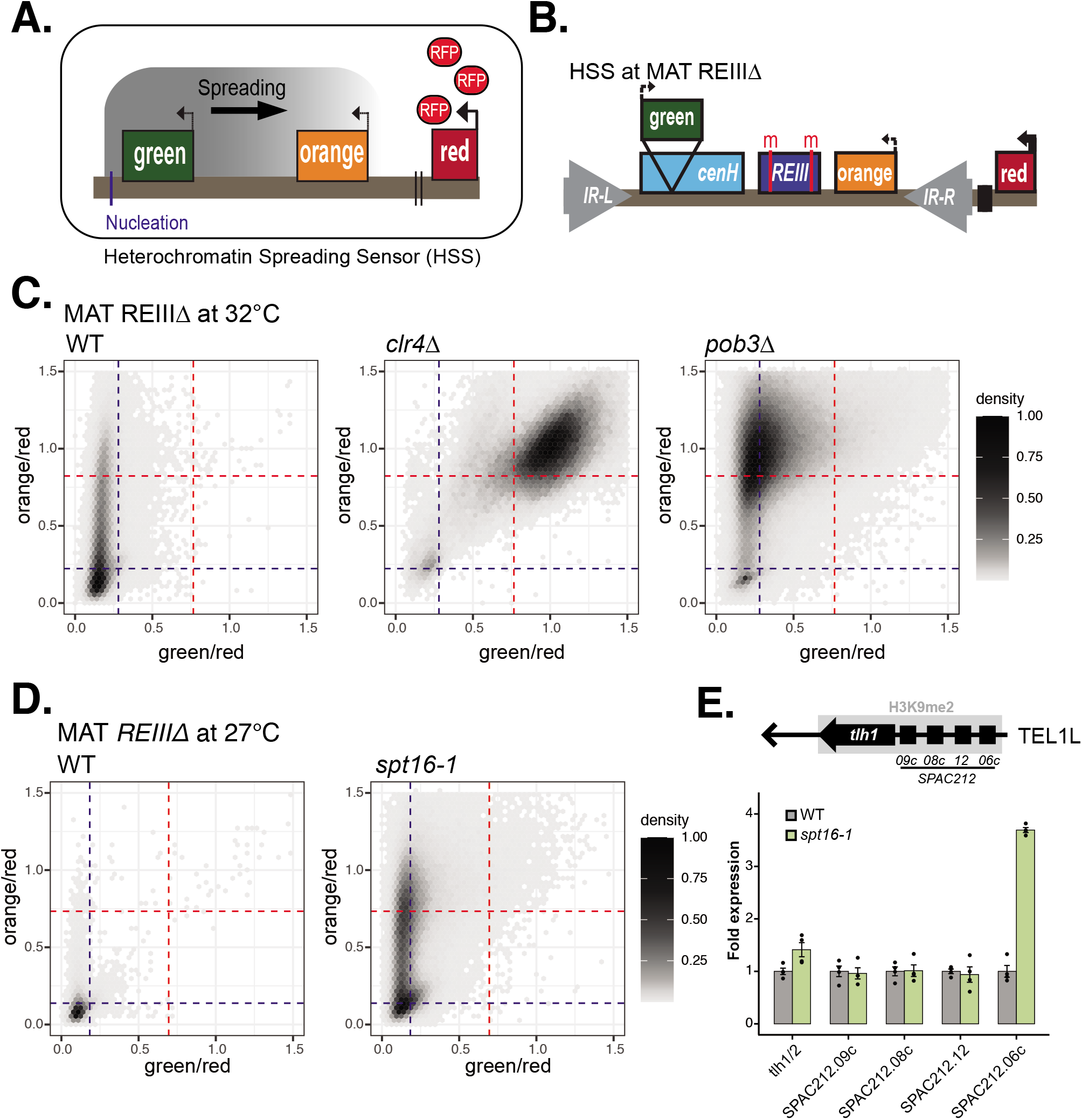
FACT mutants have heterochromatin spreading defects at engineered and endogenous heterochromatin. A. Scheme of the heterochromatin spreading sensor (HSS) assay B. REIIIΔ reporter scheme. Red bars and ‘m’ indicate mutation of the Aft1/Pcr1 binding sites. C. 2D-density hexbin plot showing the red-normalized green and orange fluorescence for WT, *clr4Δ* and *pob3Δ* MAT_REIIIΔ cells grown at 32°C. A density bar represents fraction of most dense bin. Threshold values for the fully expressed state (“on”) and fully repressed state (“off”) in each color are indicated by red and blue guide-lines respectively. One (WT, *clr4Δ*) or four (*pob3Δ*) independent isolates were analyzed and are shown in a combined 2D density hexbin plot. D. 2D-density hexbin plot showing the red-normalized green and orange fluorescence for WT and *spt16-1* MAT_REIIIΔ cells grown at 27°C. A density bar represents fraction of most dense bin. Threshold values for the fully expressed state (“on”) and fully repressed state (“off”) in each color are indicated by red and blue guide-lines respectively. One (WT) or four (*spt16-1*) independent isolates were analyzed and are shown in a combined 2D density hexbin plot. E. RT-QPCR analysis. Expression of transcripts at subtelomere 1L (TEL1L) at 27°C in the *spt16-1* strain relative to WT after normalization to *act1*+. The TEL1L gene array scheme is shown above the graph. n=4 biological replicates; data are represented as mean +/-SEM.

The partial loss-of-function of *spt16-1* allowed us to assess heterochromatin spreading in the HSS assay in this mutant. When examining spreading in the WT strain, we found that the two reporters were fully repressed in the WT strain at 27°C (Fig. 2D; compare with WT cells at 32°C in Fig. 2C). Increased silencing at lower temperature is not unexpected, as heterochromatin spreading is temperature-sensitive (Greenstein et al., 2018). In contrast, in *spt16-1* cells, the spreading reporter was partially derepressed, while the nucleation reporter remained largely repressed (Fig. 2D), revealing a heterochromatin spreading defect also in this FACT mutant.

Since H3K9me2 was reduced towards the heterochromatic boundaries at subtelomeres in *pob3Δ* (Fig. 1A), we next investigated potential spreading defects at these loci in the *spt16-1* mutant by examining the expression of several subtelomeric genes under the permissive temperature. Even though *spt16-1* does not display silencing defects at pericentromeric heterochromatin at 27°C (Fig. S2F), we observed de-repression of several subtelomeric genes (Fig. 2E and S2G). Remarkably, those genes with silencing defects are located close to telomere-distal heterochromatin at TEL1L and TEL1R, but not telomere-proximal where nucleation is mediated by shelterin and RNAi. Taken together, our results strongly suggest that FACT has a specific function in heterochromatin spreading at the mating-type and subtelomeric heterochromatin loci.

### Epe1 pathway specifically counteracts FACT at heterochromatin

To identify pathways by which FACT controls heterochromatin spreading, we sought to investigate potential suppressors of the *pob3Δ* mutant. Several mutants counteract heterochromatin stability and spreading, including the histone acetyltransferase Mst2 (Flury et al., 2017; Georgescu et al., 2020; Wang et al., 2015), the transcription elongation complex Paf1C (Kowalik et al., 2015; Sadeghi et al., 2015; Verrier et al., 2015), and the Jumonji protein Epe1, which shows homology to histone demethyltransferases and acts as heterochromatin boundary factor (Ayoub et al., 2003; Braun et al., 2011; Zofall and Grewal, 2006). We generated double mutants of those genes with *pob3Δ* and monitored silencing by growth-based silencing reporter assays and RT-QPCR (Fig. 3). Cells lacking *pob3+* showed, as expected, heterochromatin silencing defects for the *ura4+* reporter integrated at the *mat* locus (Fig. 3A, 3C and 3E). Deletions of *leo1+* or *mst2+* did not suppress *pob3Δ* silencing defects at the mating-type, pericentromeric, and subtelomeric heterochromatin (Fig. 3A–D). This result is in agreement with the predominant functions of these factors in maintaining euchromatin by preventing heterochromatin spreading or its ectopic assembly beyond its natural boundaries. In contrast, deletion of *epe1+* suppressed the *pob3Δ* silencing defects nearly to levels as seen in WT cells at all tested heterochromatic regions (Fig. 3E, 3F). Moreover, *epe1Δ* reduced the expression of several of the subtelomeric genes in the *pob3Δ* mutant, suggesting that it also counteracts heterochromatin spreading (Fig. 3G). To test this notion more directly, we performed the HSS assay in the double *spt16-1epe1Δ* mutant. Indeed, heterochromatin spreading was completely restored in *spt16-1* in the absence of Epe1 (Fig. 3H, compare with Fig. 2D). Together, our suppressor analysis revealed that the Epe1 pathway specifically suppresses silencing and spreading defects of FACT mutants, supporting the distinct function of the FACT chaperone in heterochromatin spreading.

Epe1 is recruited to heterochromatin via Swi6, which binds to H3K9me3 (Zofall and Grewal, 2006). The specific suppression of the FACT silencing and spreading defects by *epe1Δ* raised the possibility that Epe1 steady state levels or abundance at heterochromatin are increased in the FACT mutants. To test this hypothesis, we examined the turnover of Epe1, which is mediated by the Cul4-Ddb1-Cdt2 ubiquitin ligase complex during S-phase (Braun et al., 2011). Although degradation of Epe1 still occurs when Pob3 is absent, as seen for cells enriched in S phase, we found indeed increased steady-state levels of Epe1 in cycling cells (Fig. S3A). This may suggest that Epe1 is more strongly recruited to heterochromatin when FACT is impaired. However, ChIP-QPCR of Epe1 in the *pob3Δ* and *spt16-1* strains showed no increased accumulation of Epe1 at pericentromeres or subtelomeres (Fig. S3B). Instead, we found decreased levels of Epe1 at subtelomeres in *pob3Δ*, which is consistent with reduced H3K9me2 levels at those loci in this strain (Fig. 1A). Based on these findings, we reasoned that the heterochromatin spreading defect in FACT mutants is likely not due to increased Epe1 accumulation at heterochromatin. However, it is also possible that heterochromatin alterations in the FACT mutants promote transient or unstable Epe1 binding which cannot be captured by ChIP assay.

**Figure 3.**
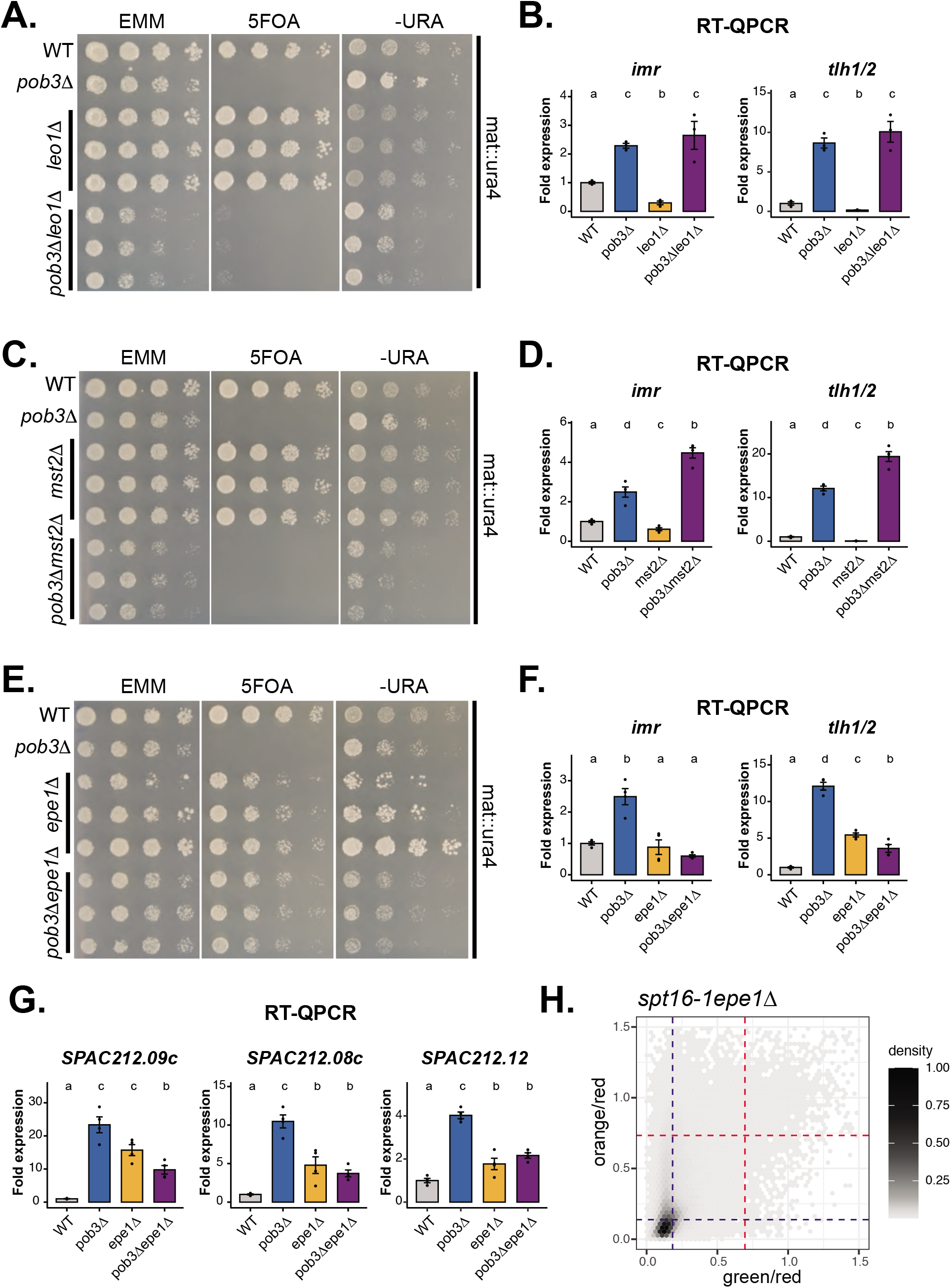
Deletion of *epe1+* suppresses FACT silencing and spreading defects. A. Silencing reporter assay at *mat* locus. Fivefold serial dilutions of WT, *pob3Δ, leo1Δ* and 3 independent *pob3Δleo1Δ* isolates were grown on the indicated media. B., D. and F. RT-QPCR analysis. Expression of *imr* and *tlh1/2* transcripts in the indicated strains relative to WT after normalization to *act1*+. n=3 biological replicates (B.) or n=4 biological replicates (D. and F.); data are represented as mean +/−SEM. Statistical analysis was done on log2 transformed values. One-way ANOVA was performed, different letters denote significant differences with a Tukey’s post hoc test at P < 0.05. C. Silencing reporter assay at *mat* locus. WT, *pob3Δ*, *mst2Δ* and 3 independent *pob3Δmst2Δ* isolates were grown on the indicated media as in A. E. Silencing reporter assay at *mat* locus. WT, *pob3Δ, epe1Δ* and 3 independent *pob3Δepe1Δ* isolates were grown on the indicated media as in A. G. RT-QPCR analysis. Expression of transcripts at subtelomere 1L (TEL1L) at 27°C in the *pob3Δ*, *epe1Δ* and *pob3Δepe1Δ* strains relative to WT after normalization to *act1*+. n=4 biological replicates; data are represented as mean +/-SEM. Statistical analysis was done on log2 transformed values. One-way ANOVA was performed, different letters denote significant differences with a Tukey’s post hoc test at P < 0.05. H. 2D-density hexbin plot showing the red-normalized green and orange fluorescence for *spt16-1epe1*Δ MAT_REIIIΔ cells grown at 27°C. A density bar represents fraction of most dense bin. Threshold values for the fully expressed state (“on”) and fully repressed state (“off”) in each color are indicated by red and blue guide-lines respectively. Three independent isolates were analyzed and are shown in a combined 2D density hexbin plot. See also Figure S3.

### FACT promotes the transition of H3K9me2 to H3K9me3 through histone turnover suppression

Having established a specific function of FACT in heterochromatin spreading, we sought to elucidate the underlying mechanism and inspected closer the heterochromatin structure in the *spt16-1* mutant. H3K9me2 and H3K9me3 methylation states have distinct heterochromatic functions. H3K9me2 domains are transcriptionally active and sufficient for RNAi-dependent co-transcriptional gene silencing (Jih et al., 2017). Conversely, H3K9me3-marked chromatin is refractory to the transcription machinery, directs RNAi-independent gene silencing, and preferentially retains Swi6 (Jih et al., 2017; Schalch et al., 2009; Yamada et al., 2005). Intriguingly, while H3K9me2 levels were unaltered in the *spt16-1* strain, H3K9me3 levels were reduced at the TEL1L subtelomeric region (Fig. 4A, 4B). This was not seen at the pericentromeres, which are mainly cotranscriptionally repressed through RNAi. This was also different from pob3Δ cells, which showed a significant decrease of H3K9me2 at subtelomeres (Fig. 1A), suggesting that this mutant displays broader phenotypes, which may be due to pleiotropic and/or indirect effects. Next, we combined a clr4^F449Y^ mutant, impaired in the transition from H3K9me2 to H3K9me3 (Jih et al., 2017), with the *spt16-1* allele, and observed a synthetic defect in the silencing of *dg* and *tlh1/2* transcripts (Fig. 4C). Since efficient heterochromatin spreading requires a self-propagating loop between H3K9 methylation by Clr4 and Swi6 binding to H3K9me, we also monitored Swi6 levels in *spt16-1*. We found a reduced Swi6 binding at the genes closer to the TEL1L heterochromatin boundary, in agreement with the spreading phenotype in this mutant (Fig. 4D). Together, our results suggest that the transition of H3K9me2 to H3K9me3 is impaired in the *spt16-1* mutant which likely impairs the heterochromatin spreading process.

**Figure 4.**
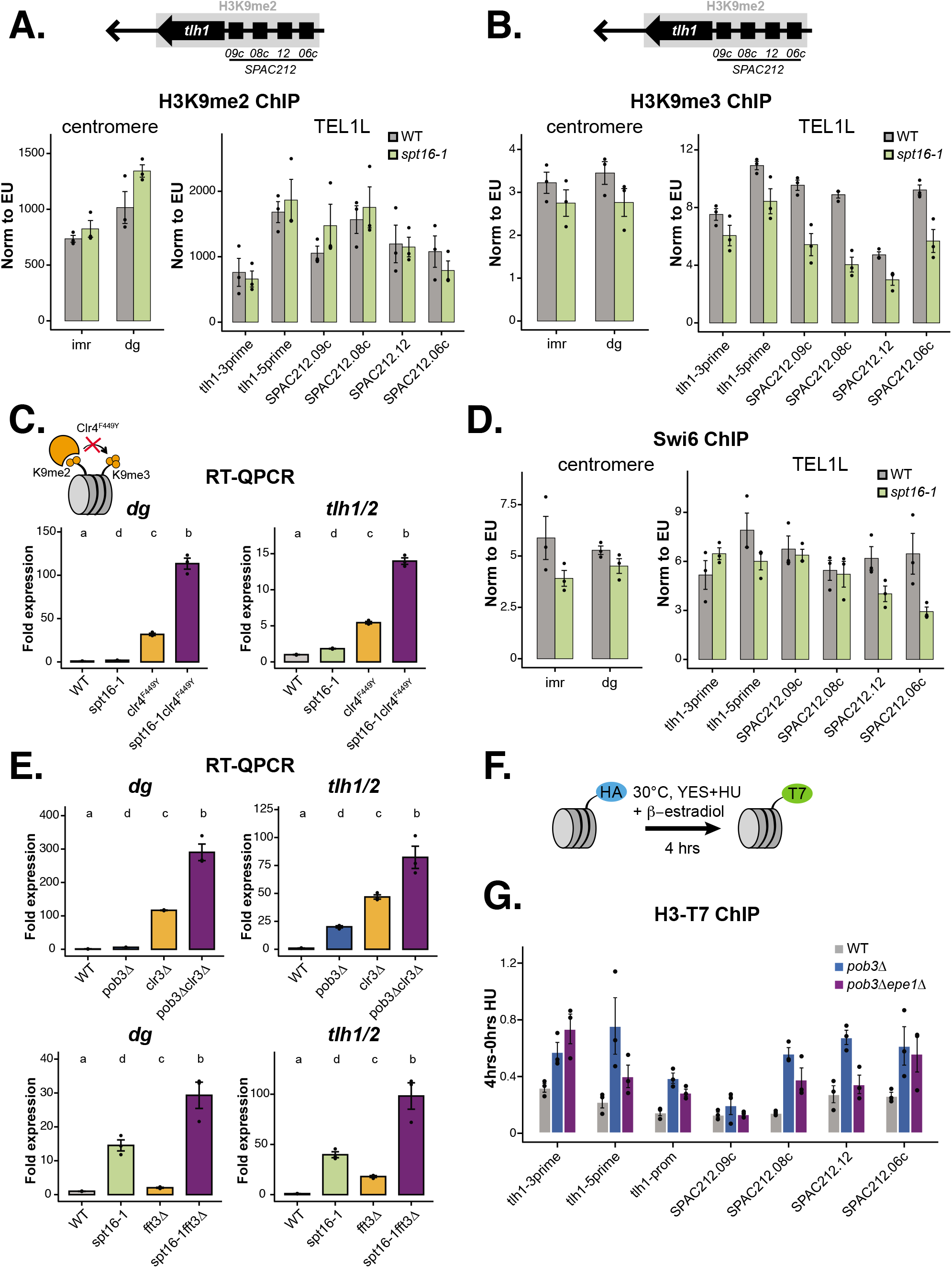
FACT facilitates H3K9me2 to H3Kme3 transition by suppressing histone turnover. A-B, D. H3K9me2 (A.), H3K9me3 (B.) and Swi6 (D.) ChIP-QPCR at pericentromere and TEL1L in the WT and *spt16-1* strains. The TEL1L gene array is shown above the graphs. ChIP was normalized to the average of tree euchromatic regions. n=3 biological replicates; data are represented as mean +/− SEM. C. RT-QPCR analysis. Expression of *dg* and *tlh1/2* transcripts in the indicated strains relative to WT after normalization to *act1*+. n=3 biological replicates; data are represented as mean +/−SEM. Statistical analysis was done on log2 transformed values. One-way ANOVA was performed, different letters denote significant differences with a Tukey’s post hoc test at P < 0.05. E. RT-QPCR analysis. Expression of *dg* and *tlh1/2* in the indicated strains relative to WT after normalization to *act1*+. The spt16-1, *fft3Δ*, *spt16-1fft3Δ* and corresponding WT were shifted to 37°C for 1.5 hour. n=3 biological replicates; data are represented as mean +/−SEM. Statistical analysis was done on log2 transformed values. One-way ANOVA was performed, different letters denote significant differences with a Tukey’s post hoc test at P < 0.05. F. Scheme of the histone turnover assay. G. ChIP QPCR of the new histone (H3-T7) at TEL1L in the WT, *pob3Δ* and *pob3Δepe1Δ* strains. Input normalized ChIP signals from the uninduced samples (0 hr) were subtracted from the input normalized signals from the *β*-estradiol-induced samples (4 hrs). Error bars represent +/− SEM from 3 independent experiments. See also Figure S4.

How can a histone chaperone facilitate the transition of H3K9me2 to H3K9me3? In contrast to mono- and dimethylation, methylation rate of H3K9me3 by Clr4 is very slow (Al-Sady et al., 2013). The molecular mechanism of this methylation step by Clr4 is not fully understood. A recent study suggested that histone turnover rates are critical for establishing methylation states of the genome (Chory et al., 2019). Noteworthy, an increased histone turnover was recently shown in *pob3Δ* for the mating-type locus (Holla et al., 2020). Thus, we hypothesized that increased histone turnover at heterochromatin in the absence of FACT impairs H3K9 tri-methylation by Clr4. Indeed, we found that both, *pob3Δ* and *spt16-1* strains, showed synthetic genetic interactions with two other factors known to repress histone turnover at heterochromatin, the histone deacetylase Clr3 (Aygun et al., 2013) and the chromatin remodeler Fft3 (Taneja et al., 2017) (Fig. 4E). Histone H3 ChIP-seq revealed a small but reproducible reduction of H3 at subtelomeres in the *pob3Δ* strain (Fig. S4A, S4B and S4C). This subtle histone change may indicate an increased nucleosome instability due to elevated histone turnover. To further test our hypothesis, we used the Recombination Induced Tag Exchange (RITE) approach (Svensson et al., 2015) to monitor incorporation of new H3 histones marked with a T7 epitope at subtelomeric heterochromatin in the *pob3Δ* cells (Fig. 4F). Both, WT and *pob3Δ* strains displayed similar tag switch recombination rates (76.5% for WT vs. 73.4% for *pob3Δ*) and H3-T7 signals were not increased at a control euchromatic region (Fig. S4D), Nonetheless, the *pob3Δ* mutant displayed increased incorporation of T7-tagged H3 histones at the TEL1L region (Fig. 4G), implying that the H3 turnover rate is increased at subtelomeric heterochromatin.

Since Epe1 was implicated before in promoting histone turnover (Aygun et al., 2013), we tested whether *epe1+* deletion suppresses the elevated histone turnover in the FACT mutant. Indeed, using the RITE approach, we found that histone turnover at the TEL1L region (but not at pericentromers) was decreased in the double *pob3Δepe1Δ* mutant compared to *pob3Δ* (Fig. 4G and S4D). Notably, the restored histone stability in the *pob3Δepe1Δ* mutant correlated with the reduced RNAPII levels at the subtelomeric region in this strain (Fig. S4E). We conclude that the specific suppression of heterochromatin spreading by *epe1Δ* in the FACT mutants is related to the role of Epe1 in nucleosome stability regulation, in agreement with a recent study (Raiymbek et al., 2020). Altogether, our data indicate that FACT regulates heterochromatin spreading by promoting the transition of H3K9me2 to H3K9me3 through histone turnover suppression.

## Discussion

The involvement of FACT in heterochromatin silencing has been known for a long time, however the underlying mechanisms have been unclear. In addition to silencing defects caused by the absence of FACT (Holla et al., 2020; Lejeune et al., 2007), FACT has been co-purified with HP1 complexes, suggesting a direct function in heterochromatin regulation (Fischer et al., 2009; Motamedi et al., 2008). One limiting factor of previous studies was the usage of the *pob3Δ* strain, which due to its strong silencing and pleiotropic effects, likely masked specific roles of FACT in heterochromatin formation process. Here, by using a partial loss of function, *spt16-1* mutant, we were able to show that FACT promotes heterochromatin spreading process. First, we show at single-cell level using the HSS assay that the spreading reporter is derepressed while the nucleation reporter remains largely silenced in both the *pob3Δ* and *spt16-1* mutants. Second, genes distal from nucleation sites but proximal to the subtelomeric heterochromatin boundaries are de-repressed in the *spt16-1* mutant. Third, H3K9me3 and Swi6 binding, but not H3K9me2, are reduced in the spreading-deficient *spt16-1* strain at subtelomeric heterochromatin. Fourth, FACT acts synergistically with the Clr4 me2-me3 transition mutant. Finally, deletion of *epe1+*, which counteracts heterochromatin spreading, suppresses most of the silencing and spreading defects of the FACT mutants and this suppression is linked to the function of Epe1 in stabilizing nucleosomes. Based on our results, we propose that FACT facilitates heterochromatin spreading through maintaining low level of histone turnover which enables a productive H3K9 tri-methylation step by Clr4.

Our analysis revealed also important differences between the two FACT mutants. While H3K9me2 levels are gradually reduced from the subtelomeric nucleation sites in *pob3Δ*, H3K9me2 is not changed in the *spt16-1* mutant. In contrast to *pob3Δ*, which has strong silencing defects genome-wide, *spt16-1* shows de-repression of genes in close proximity to the heterochromatin boundaries, in agreement with its spreading defects. Thus, our results highlight the importance of hypomorphic mutants for studying abundant and pleiotropic protein complexes.

Broadly, our results link several studies that implied H3K9me3 function in heterochromatin spreading (Al-Sady et al., 2013; Jih et al., 2017). They are in agreement with the idea that histone turnover rate is a crucial factor that determines methylation states of the genome (Becker et al., 2016; Chory et al., 2019). Particularly the low histone turnover may be important for enzymes with slow kinetic rates, like Clr4, which catalyzes K9me2 - me3 roughly ten times slower than the mono- and di-methylation steps (Al-Sady et al., 2013). Similar slow kinetic properties are shared for human K9 and K27 methyltransferases, G9a (Patnaik et al., 2004) and EZH2 (Alabert et al., 2020; Chory et al., 2019), respectively. Thus, low histone turnover emerges as a critical factor for the establishment of repressive chromatin domains.

Histone turnover was recently linked to heterochromatin inheritance and epigenetic memory (Aygun et al., 2013; Greenstein et al., 2018; Holla et al., 2020; Shan et al., 2020; Taneja et al., 2017). Our results support the findings that histone chaperones which guide heterochromatic histone turnover may be broadly involved in controlling cell fate (Brumbaugh et al., 2019; Cheloufi et al., 2015; Kolundzic et al., 2018). Future studies should address whether FACT function in heterochromatin spreading control is conserved in metazoan development and cell fate maintenance.

## Acknowledgments

We thank LAFUGA and Stefan Krebs and Helmut Blum for sequencing. We thank Julia Schluckebier for assistance in strain generation. This research was supported by the European Commission (Marie-Curie Individual Fellowship H2020-MSCA-IF-2014 contract 657244 to M.M.; Network of Excellence EpiGeneSys HEALTH-2010-257082 to S.B.), the Friedrich-Baur-Stiftung (to M.M. and S.B.) and DFG (project number 453441129 to M.M.), grants from the National Institutes of Health (DP2GM123484 to B.A-S.) and the UCSF Program for Breakthrough Biomedical Research (to B.A-S.).

## Author Contributions

Conceptualization, M.M., S.B, B. A-S. and A.G.L.; Investigation, M.M., R.A.G., K.C.F.O., H.N., B. A-S.; Formal Analysis, M.M., T.S., R.A.G.; Visualization, M.M., T. S., R.A.G. and B. A-S; Resources, S.B., B. A-S., and A.G.L.; Writing – Original Draft, M.M., Writing – Review & Editing, M.M., S.B., B. A-S. and A.G.L., Funding acquisition, M.M., S.B., B. A-S and A.G.L.

## Declaration of Interests

The authors declare a competing interest. A.G.L. is co-founder and CSO of Eisbach Bio GmbH.

## Material and methods

### Yeast strains, media, and growth conditions

*S. pombe* strains used in this study are listed in Table S1. Strains were generated with standard procedures using yeast transformation and validated by colony PCR. Point mutants were generated using a CRISPR/Cas9 system according to the published method (Torres-Garcia et al., 2020). All CRISPR/Cas9 generated strains were sequenced to confirm the presence of the mutation. Strains used for RITE assay were generated by crossing out the cdc25-22 allele using random spore analysis. Strains were grown in rich media (YES) at 32°C, 30°C or at 27°C as indicated. For temperature sensitive alleles, strains were grown at 26°C overnight and then they were shifted to 36°C for 1.5 hours. 5-FOA medium contained 1 g/L 5′-fluoroorotic acid.

### RNA extraction and cDNA synthesis

RNA extraction and gene expression analysis were done as described in (Barrales et al., 2016; Murawska et al., 2020). Briefly, 50 mL of yeast culture at OD600 0.5-0.8 was spun down at RT and the pellet was frozen in liquid nitrogen. Cells were thawed on ice and resuspended in 1 mL of TRIzol. 250 μl of zirconia beads were added and cells were broken with Precyllis 24 (Peqlab) for 3×30 sec with 5 min rest on ice. The extract was spun down at 13500 rpm at 4°C for 10 min. The cleared lysate was extracted twice with chloroform and spun at 13500 rpm at 4°C for 10 min. The aqueous phase was taken and RNA was precipitated with isopropanol. The pellet was washed twice with 75% EtOH, air-dried and resuspended in 50 μL of RNase free dH20. The RNA concentration and purity were determined by Nano-drop. For RT-QPCR 20 mg of RNA was treated with 1 μL of TURBO DNase I (Ambion) for 1 hr at 37°C. The reaction was inactivated by adding 6 μL of DNase inactivation reagent followed by the manufacturer instructions. For cDNA synthesis 5 μg of total DNase-treated RNA was reverse transcribed with 1 μL of oligo-(dT)20 primers (50 μM) and 0.25 μL of SuperscriptIII (Invitrogen) according to the manufacturer instructions.

### Gene expression analysis

cDNAs were quantified by qPCR using Fast SYBR Green Master mix (Life Technologies) and a 7500 Fast real-time PCR system (Applied Biosystems). cDNA was analyzed by qPCR using gene specific primers (Table S2). The quantification was based on a standard curve method obtained with QuantStudioTM Design and Analysis Software. Sheared *S. pombe* genomic DNA was used as a standard. For gene expression the samples were normalized to *act1* gene. The normalized datasets were shown as relative to the mean value of the WT strain which was set to 1, errors bars were calculated as SEM and displayed accordingly.

### ChIP-seq and ChIP-QPCR

ChIP-QPCR and ChIP-seq were performed as described previously (Murawska et al., 2020). Briefly, 100 mL yeast cultures were grown to mid-log phase to OD600 = 0.6. The cultures were cooled down at RT for 10 min and fixed with 1% FA for 20 min at RT on the shaker. Cross-linking was stopped with 150 mM Glycine for 10 min at RT. Cells were washed 2x with 30 mL ice-cold dH2O, the pellet was frozen in liquid nitrogen and kept at −80°C until further processing. Pellets were resuspended in 1 mL FA(1) buffer (50 mM HEPES-KOH, pH 7.5, 150 mM NaCl, 1 mM EDTA, 1% Triton X-100 (v/v), 0,1% NaDeoxycholate (w/v), 0,1% SDS (w/v), supplemented with Roche protease inhibitors. Cells were broken in a bead beater (Precellys): 9×30s (for ChIP-seq or 6×30s for ChIP-QPCR). After centrifugation, the supernatant and the pellet were sonicated for 15 min at 4°C. Chromatin extracts were spun down for 10 min at 14000 rpm at 4°C. Different amount of chromatin was used for different antibodies: 100 μl chromatin (for H3-T7, Spt16, Pob3 ChIP) or 500 μl chromatin (for Pol IISer2, H3K9me2, H3K9me3, H2Bub, Swi6, Clr4-HA, Epe1-Flag ChIP). The antibodies used for ChIP are listed in Key Resources Table. Samples were incubated with antibodies O/N at 4°C. 25 μl of FA(1) buffer washed Dynabeads were added to each sample and they were incubated for 2 hours at 4°C. Samples were then washed 3x for 5 min at RT with FA(1) buffer, FA(2) buffer (FA(1) buffer with 500 mM NaCl), once with LiCl buffer (10 mM TrisHCl, pH 8.0, 0.25 M LiCl, 1 mM EDTA, 0,5% NP-40 (v/v), 0,5% NaDeoxycholate (w/v)) and once with TE buffer. DNA was eluted from the antibodies with ChIP Elution buffer (50 mM Tris HCl, pH 7.5, 10 mM EDTA, 1% SDS) for 15 min at 65°C in a thermomixer set to 1300 rpm. DNA was treated with Proteinase K and de-crosslinked O/N at 65°C. DNA was purified with Zymo Research ChIP DNA Clean and Concentrator kit according to the manual instructions. To obtain enough material for the library preparation usually 3 technical IP replicates were pulled. The ChIP-seq libraries were prepared with 2 ng of DNA with NEBNext Ultra II DNA Library Prep Kit for Illumina according to the manual instructions. The libraries were barcoded and sequenced at LAFUGA at the Gene Center (LMU). For ChIP-QPCR, the isolated DNA was quantified by qPCR as described for the gene expression analysis. Unless otherwise noted, the mean was calculated from three independent experiments and errors bars were calculated as SEM and displayed accordingly. QPCR signals were normalized against the input samples for each primer position as internal control. For ChIP experiments with anti-H3K9me2, anti-Swi6, anti-H3K9me3, the input normalized values were corrected for variation in IP efficiency by normalizing against the mean of 3 euchromatin loci (act1, adh1-prom, adh1-5prime). For RNAPII Ser2P ChIP, the input normalized values were corrected for variation in IP efficiency by normalizing against the mean of 3 euchromatin loci (act1, ade2, adh1-3prime). Euchromatin normalized ChIPs were displayed as ‘Norm to EU’. For ChIP experiments with anti-Pob3, anti-Spt16 and anti-HA (Clr4-HA ChIP) ChIP signals were normalized to input and displayed as ‘% of Input’. For the histone turnover ChIP, incorporation of the new histone H3-T7 was calculated as follows: input normalized signals from the β-estradiol-uninduced samples were subtracted from the input normalized signals from the β-estradiol-induced samples as displayed as ‘4hrs-0hrs’ HU.

### ChIP-seq analysis

Single-end reads (50 bp) were mapped to the reference genome (Schizosaccharomyces pombe ASM294v2) using bowtie2 (version 2.2.9).

Reads were counted in either 250 bp or 10 kb windows (bins) using the windowCounts() function from the csaw R/Bioconductor package (version 1.18.0). Normalization factors were calculated by the normFactors() function using the “TMM” method on count matrices with 10 kb bins and applied on count matrices with 250 bp bins. Normalized, log2-transformed count matrices were obtained by the cpm() function (edgeR package, version 3.26.8). ChIP samples were normalized by their corresponding inputs (i.e. subtraction in log2-scale). Input-normalized count matrices were converted to coverage vectors using the coverage() function (GenomicRanges package, version 1.38.0) and exported as bigwig files (rtracklayer package, version 1.44.4). For visualization, coverages were smoothed by the rollmean() function (zoo package, version 1.8.9) and plotted using custom R functions.

Broad H3K9me2 peaks were identified using the Homer software package (Heinz et al., 2010). Tag directories were created with the settings -mapq 1 and parameters for findPeaks command were set to -style histone -F 2. Peaks were called on replicates independently and the intersect was taken as final set.

Normalized count matrices (with 250 bp bins) were subset by telomeric, centromeric heterochromatin or euchromatin regions on chromosome I and II. Telomeric and centromeric bins were selected by the overlap of wild-type H3K9me2 peaks with telomeres (I:1-100000; I:5479000-5579000; II:1-100000; II:4440000-4539000) or centromeres (I:3720000-3820000; II:1570000-1670000), respectively. The data were aggregated using two approaches: either replicates were averaged and values of bins were visualized as boxplots or the median of bins were calculated for each replicate and region, and visualized as dot plots. Statistical analysis was performed on aggregated data using the second approach (i.e. on medians). P-values were obtained by fitting a linear mixed effect model (lme4,_version 1.1-27 and lmerTest, version 3.1-3 packages) with genotype (e.g. wild-type or *pob3Δ*) and region (e.g. telomeric, centromeric or euchromatin) as fixed effects and sample id as random intercept.

### Histone turnover assay

RITE histone turnover assay was done as before (Greenstein et al., 2018) with several modifications. Briefly, cells were inoculated from a pre-culture to 100 ml YES supplemented with Hygromycin B (100 μg/ mL, Invitrogen) and grown O/N at 30°C to OD600 0.4-0.8. 20 ODs of cells were taken and processed for ChIP as the 0 hr (uninduced) time point. The remaining cells were washed 2x in media devoid of Hygromycin B. 12.5 ODs of cells were taken and resuspended in 50ml YES supplemented with 15 mM Hydroxyurea (HU) and 1.5 μM β-Estradiol (ER) and incubated further for 4 additional hours at 30°C. 20 ODs of cells were processed for ChIP as the 4 hrs (induced) time point.

### Protein extract preparation

Whole cell extracts (WCE) were prepared as published before (Murawska et al., 2020). Briefly, 50 mL yeast cultures were grown to mid-log phase (OD600 0.5-0.8). The cultures were spun down and cell pellets were resuspended in 500 μL of Workman Extract Buffer (40mM HEPEs pH7.4, 250mM NaCl, 0.1% NP40, 10% Glycerol, 1 mM PMSF, Roche proteinase inhibitors). 250 μL of glass beads were added and cells were lysed with Peqlab precellys homogenizator (3×30 sec). The extracts were shortly spun down at 2500 rpm at 4°C. The supernatant and the pellet were treated with benzonase in the presence of 2mM MgCl2 for 30 min on ice and spun down at the maximum speed for 10 min at 4°C. The extracts were frozen in liquid nitrogen and stored at −80°C or immediately used for Western blot analysis. Protein concentration was measured with Bradford reagent (BioRad).

For HU treatment assay, strains were pre-cultured in liquid YES at 27°C for 24 hours, back diluted and grown to log-phase. A control (untreated) culture and treated culture (20mM HU) were incubated for a further 2 hours at 27°C. Total protein extracts from OD600=1 were prepared by trichloroacetic acid (TCA) precipitation according to (Knop et al., 1999). Proteins solubilized in HU buffer (200 mM phosphate buffer, pH 6.8, 8 M urea, 5% w/v SDS, 1 mM EDTA,100 mM DTT) and heat denatured at 65°C for 10 minutes were separated with SDS-polyacrylamide gel electrophoresis and subjected to Western Blot analysis.

### Western Blot

Western blot was performed as published before (Murawska et al., 2020). Briefly, proteins were separated with SDS-polyacrylamide gel electrophoresis and electroblotted onto methanol activated polyvinylidene difluoride (PVDF) membranes in Blotting Buffer (20 mM Tris, 192 mM glycin, 20% methanol) for 1hr at 400 mA at 4°C. Membranes were then incubated in Blocking Buffer (TBS, 0.1% Tween 20, 5% non-fat dry milk) for 40 min - 1hr at RT followed by an incubation in the Blocking Buffer with the appropriate primary antibody for 1hr at RT. Membranes were washed three times for 5 min in Washing Buffer - TBST (TBS, 0.1% Tween 20) and then incubated in Blocking Buffer containing the appropriate fluorescence or HRP-conjugated secondary antibodies for 40 min - 1 hour at RT followed by 3 times washing in TBST for 5-10 min. Fluorescent Western blot signals were visualized with Li-Cor Imaging System. Chemiluminescent Western blot signals were visualized with the Immobilon Western Chemiluminescence HRP substrate (Millipore, WBKLS0500) using BioRad ChemicDoc™ MP Imaging System.

### HSS assay

Cells containing HSS reporters were grown for flow cytometry experiments as described (Greenstein et al., 2020). Flow cytometry was performed using a Fortessa X20 dual machine (Becton Dickinson) and high-throughput sampler (HTS) module. Depending on strain growth and sample volume, data from approximately 4,000–100,000 cells were collected. Fluorescence detection, compensation, and data analysis were done as described (Al-Sady et al., 2016; Greenstein and Al-Sady, 2019; Greenstein et al., 2018). 2D-density histogram plots were generated as described previously (Greenstein and Al-Sady, 2019; Greenstein et al., 2018) with the following exceptions: Data from biological replicates were merged together prior to plotting. Hexbin plots were generated in R via the ggplot2 package. The guidelines for cutoff values of “off” and “on” states for Green and Orange were determined using mean of a Red-Only control strain plus 3 times the standard deviation (SD) and mean of *clr4Δ* (after removal of color-negative cells) minus 1 SD value respectively. For the *spt16-1* experiment, the fraction of cells below the “off” threshold for both “green” and “orange” were calculated for each biological replicate independently.

### Cell cycle analysis

PAS99 wildtype, *pob3Δ::KAN* and *spt16-1:KAN* cells were struck onto YS plates, and grown at 27°C in EMM liquid culture. Cell were diluted to OD 0.1 and grown another 9 hrs in EMM at 27°C till OD ~0.5-0.8. For the *cdc25* experiment, cells were grown at 25°C overnight, diluted as above and then either kept at 25°C or moved to 37°C for 2 or 4hrs. Cells were then fixed, RNaseA treated and stained with Sytox Green as described (Knutsen et al., 2011) but instead of sonication, cells were vigorously vortexed just prior to Flow cytometry analysis. Flow cytometry and analysis was performed as described (Greenstein et al., 2018). Within each experiment, the same gates were applied to all specimen for forward and side scatter, to select a population of similar sized and shaped cells to analyze, and for Width and Area of SyTOX green fluorescence, to assess cell cycle stage distributions.

### Quantification and statistical analysis

Quantification, number of replicates and statistical tests employed are described in the figure legends or in the method section. Shortly, for RT-QPCR statistical analysis was done on log2 transformed values. One-way ANOVA was performed, and different letters were assigned with significant differences with a Tukey’s post hoc test at P < 0.05.

### Data and code availability

ChIP-seq data for H2Bub, H3, Pob3, RNAPII Ser2P and Spt16 in wild-type and pob3Δ strains were previously deposited to GEO with accession number GSE135766. H3K9me2 ChIP-seq data in wild-type and *pob3Δ* strains were deposited to GEO with accession number GSE174641.

Code for the data analysis is available on github at: https://github.com/tschauer/Murawska_etal_2021

---

**Figure S1.**
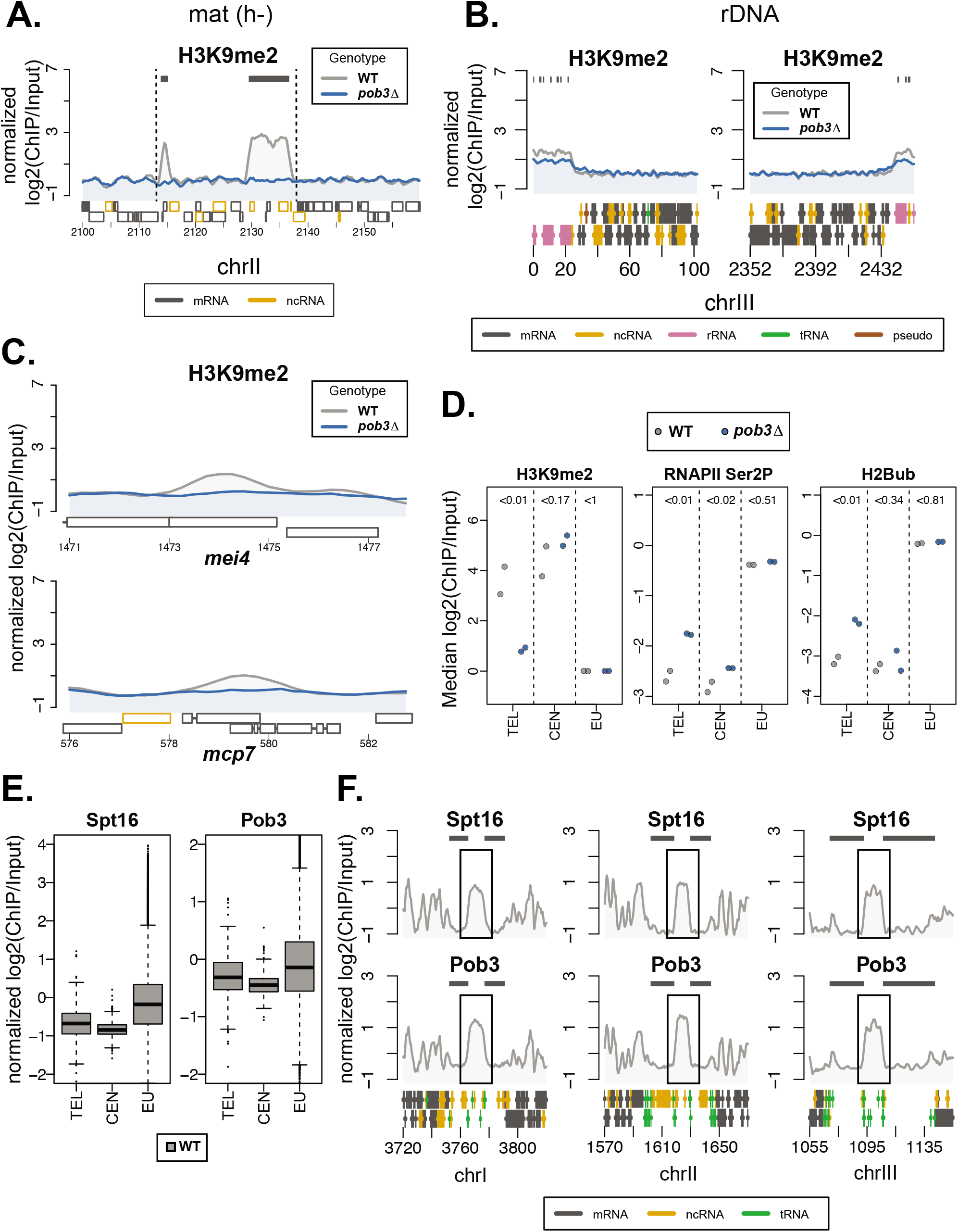
H3K9me2 level reduction in the FACT mutant at mating-type, rDNA and heterochromatic islands and analysis of FACT binding to heterochromatin regions in *S. pombe*. Related to Figure 1. A. H3K9me2 ChIP-seq enrichment [normalized log2(ChIP/Input)] at mating-type (h-) locus in WT and *pob3Δ* strains. Dark grey bars indicate H3K9me2 heterochromatic regions. Black dashed lines indicate the mating-type locus boundaries. Note the REII-cenH part of the locus is missing due to h-type of the mating-type in the analyzed strains. Average of 2 biological replicates is shown. B. H3K9me2 ChIP-seq enrichment [normalized log2(ChIP/Input)] at ribosomal genes in WT and *pob3Δ* strains. Dark grey bars indicate H3K9me2 heterochromatic regions. Average of 2 biological replicates is shown. C. H3K9me2 ChIP-seq enrichment [normalized log2(ChIP/Input)] at two examples of heterochromatin islands in WT and *pob3Δ* strains. Average of 2 biological replicates is shown. D. Median ChIP-seq enrichment [normalized log2(ChIP/Input)] of H3K9me2, RNAPII Ser2P and H2Bub in WT and *pob3Δ* strains. TEL – subtelomeres, CEN – pericentromeres, EU – rest of the genome (euchromatin). Two biological replicates are shown for each region. Statistical analysis was performed by using a linear mixed effect model, P values are indicated above each region. E. Boxplots of Spt16 and Pob3 ChIP-seq enrichment [normalized log2(ChIP/Input)] calculated in 250 bp bins in the WT strain at TEL, CEN and EU regions (labelling as in D.). Average of 2 biological replicates is shown. F. ChIP-seq enrichment [normalized log2(ChIP/Input)] of Pob3 and Spt16 at centromeres of three *S. pombe* chromosomes in the WT strain. Dark grey bars indicate H3K9me2 heterochromatic regions. Black boxes indicate the centromere cores. Average of 2 biological replicates is shown.

**Figure S2.**
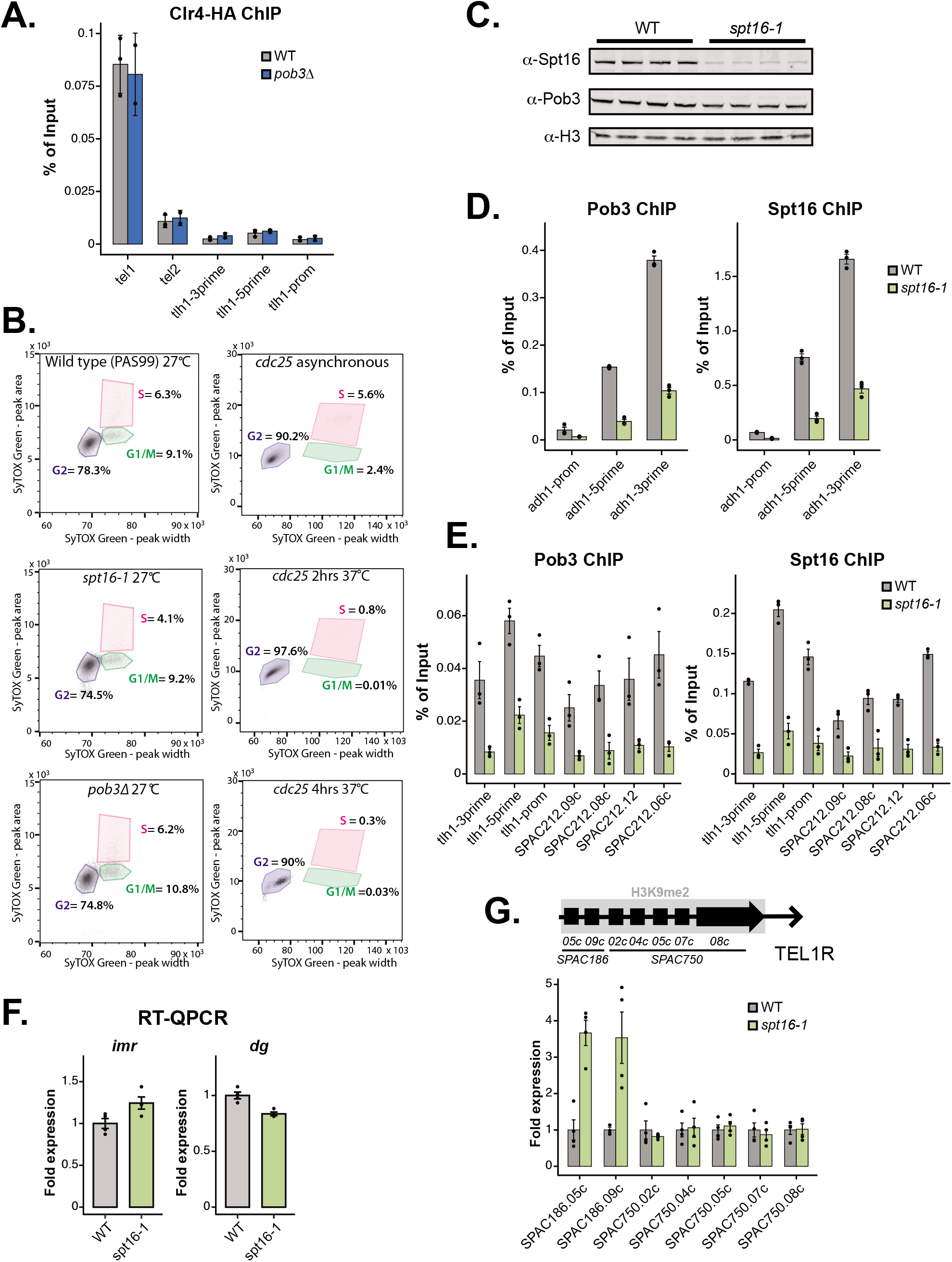
Characterization of the *spt16-1* mutant. Related to Figure 2. A. ChIP-QPCR of Clr4-HA at TEL1L region in the WT and *pob3Δ* strains. ChIP signals were normalized to input. Error bars represent +/−SD from 2-3 independent experiments. B. Cell cycle analysis of the WT, *pob3Δ* and *spt16-1* mutants at 27°C and *cdc25 ts* mutant shifted for 2 or 4 hours at 37°C, as indicated. C. Western blot for Spt16 and Pob3 protein levels in whole cell extracts from WT and *spt16-1* cultures grown at 27°C. Extracts from four cultures were loaded on the gel. Histone H3 was used as a loading control. D. ChIP-QPCR of FACT subunits, Spt16 and Pob3, at an active *adh1+* gene. Cells were grown at 27°C. ChIP signals were normalized to input. n=3 biological replicates; data are represented as mean +/−SEM. E. ChIP-QPCR of FACT subunits, Spt16 and Pob3, at TEL1L region. Cells were grown at 27°C. ChIP signals were normalized to input. n=3 biological replicates; data are represented as mean +/−SEM. F. RT-QPCR analysis. Expression of *imr* and *dg* transcripts in the *spt16-1* strain relative to WT after normalization to *act1*+. Cells were grown at 27°C. n=4 biological replicates; data are represented as mean +/−SEM. G. RT-QPCR analysis. Expression of transcripts at TEL1R region in the *spt16-1* strain relative to WT normalized to *act1*+. Cells were grown at 27°C. n=4 biological replicates; data are represented as mean +/−SEM.

**Figure S3.**
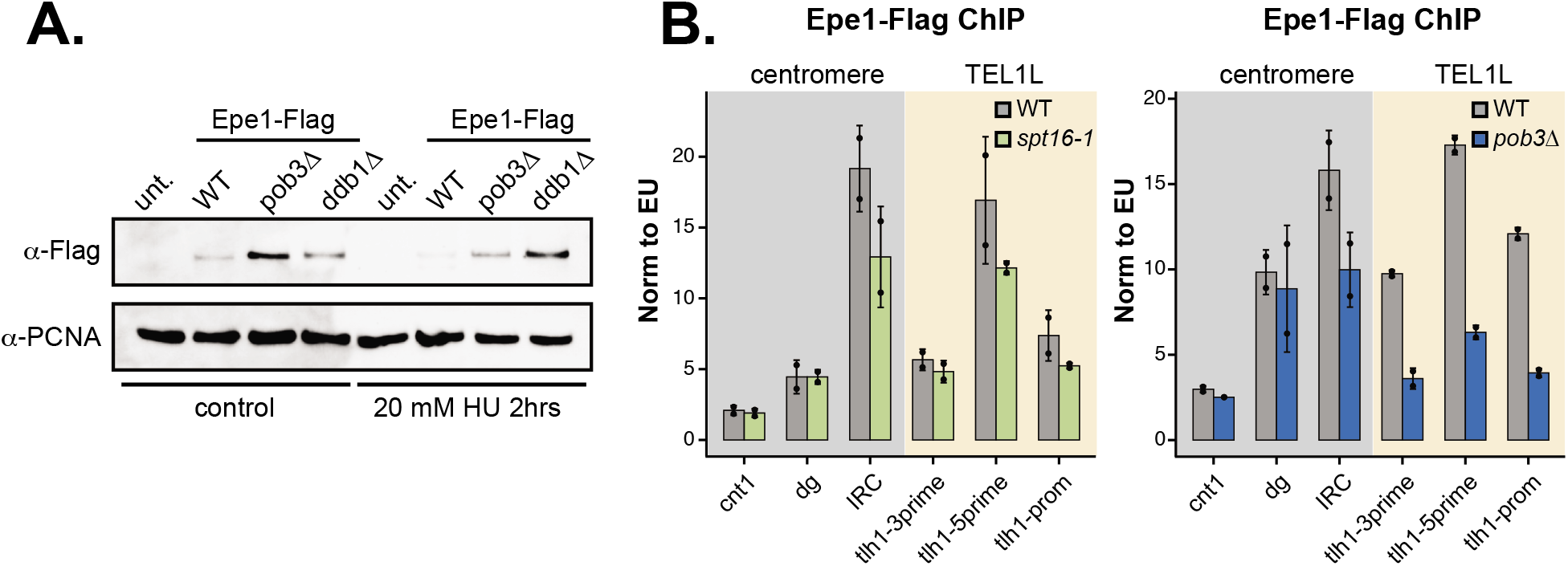
Epe1 degradation and heterochromatin binding in the FACT mutants. Related to Figure 3. A. A representative Western blot for Epe1-Flag protein levels in whole cell extracts from the indicated strains. *Ddb1Δ* was used as a control for the assay. Cultures were grown at 27°C +/− 20 mM hydroxyurea (HU) as indicated. PCNA was used as a loading control. B. ChIP-QPCR of Epe1-Flag at cnt (core centromere), *dg* (pericentromeric heterochromatin), *IRC* (pericentromeric heterochromatin boundary, a control region for Epe1 ChIP) and TEL1L regions in the WT and *spt16-1* (left panel) or in the WT and *pob3Δ* (right panel) strains. ChIP signals were normalized to the average of 3 euchromatic regions. Error bars represent +/− SD from 2 independent experiments.

**Figure S4.**
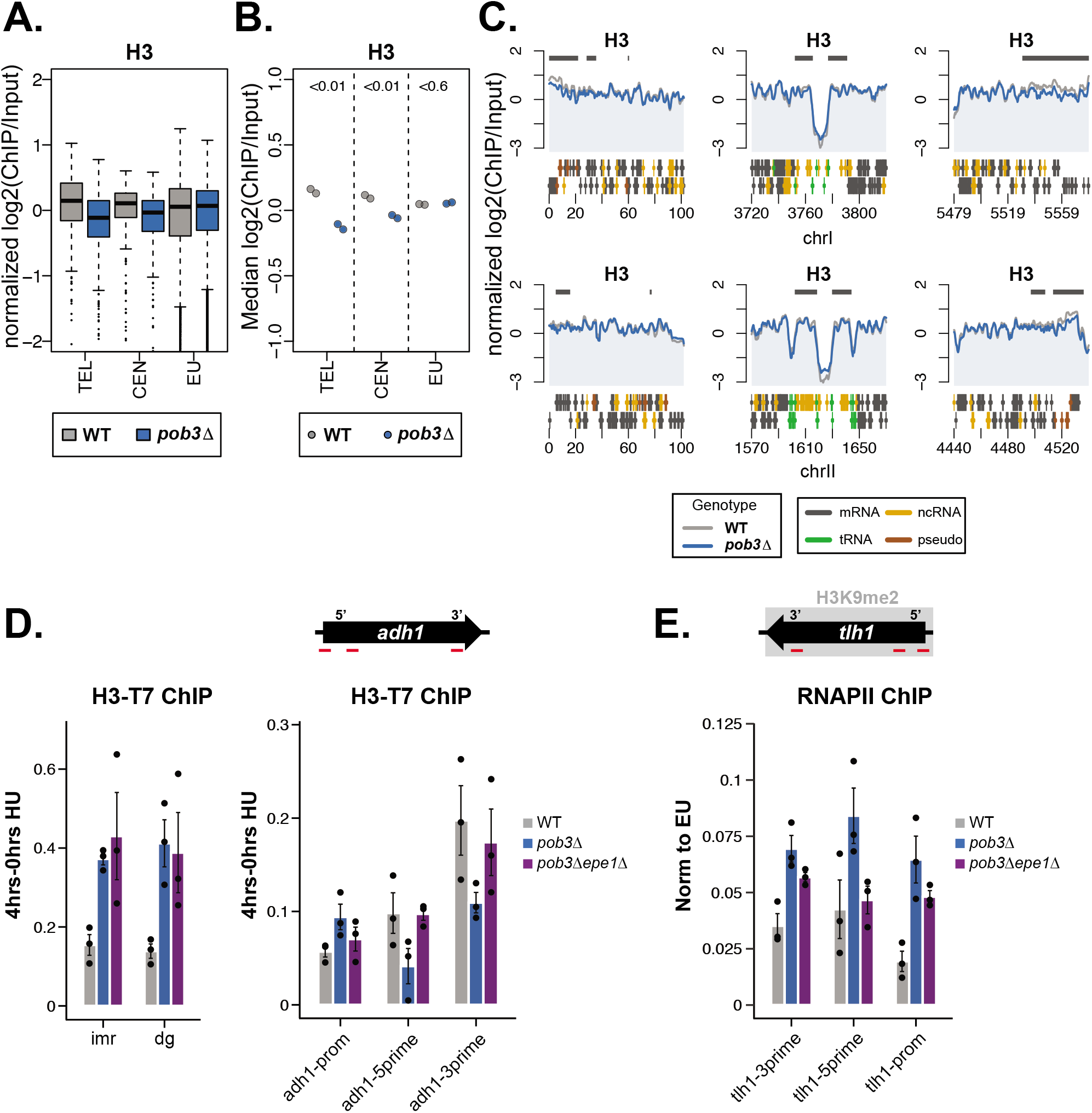
FACT maintains histone stability and low turnover at heterochromatin. Related to Figure 4. A. Boxplots of H3K9me2 ChIP-seq enrichment [normalized log2(ChIP/Input)] calculated in 250 bp bins in WT and *pob3Δ* strains. TEL – subtelomeres, CEN – pericentromeres, EU – rest of the genome (euchromatin). Average of 2 biological replicates is shown. B. Median ChIP enrichment [normalized log2(ChIP/Input)] of H3 in WT and *pob3Δ* strains at TEL, CEN and EU regions (labelling as in A.). Two biological replicates are shown for each region. Statistical analysis was performed by using a linear mixed effect model, P values are indicated above each region. C. H3 ChIP-seq enrichment [normalized log2(ChIP/Input)] at subtelomeres and pericentromeres at chromosomes I and II in WT and *pob3Δ* strains. Dark grey bars over the graphs indicate the localization of H3K9me2 in the WT strain. D. RITE histone turnover assay at pericentromere (*imr*, *dg*) and on an euchromatic gene (*adh1*) in the WT, *pob3Δ* and *pob3Δepe1Δ* strains. Primers locations (not-to-scale) are shown in the schemes above the graphs. Input normalized ChIP signals from the uninduced samples (0 hr) were subtracted from the input normalized signals from the βestradiol-induced samples (4 hrs). Error bars represent +/− SEM from 3 independent experiments. E. RNAPII ChIP-QPCR at *tlh1* gene in the WT, *pob3Δ and pob3Δepe1Δ* strains. ChIP was normalized to the average of three euchromatic regions. Error bars represent +/− SEM from 3 independent experiments.

**Table S1.** Yeast strains used in this study

| Identifier | Genotype | Figure; experiment | Source |
| --- | --- | --- | --- |
| PMM201 | h-, ade6-M216, htb1-FLAG:KANMX | Fig. 1A,1B, 1D, S1A, S1A-D; ChIP-seq | Jasson Tanny (JTB62-1) |
| PMM202 | h-, ade6-M216, htb1-FLAG:KANMX, pob3::NAT | Fig. 1A, 1B, 1D, 1E, 1F, S1A, S1B; ChIP-seq | (Murawska et al. 2020) |
| PMM01 | h-, leu1-32 ade6-210 ura4-D18 imr1L(Ncol)::ura4+ otr1R(SphI)::ade6+ mat1_m-cyhS smt0 rpl42::cyhR(sP56Q) cen1:hphMX | Fig. 1C, 1E, 1F; RT-QPCR | Sigurd Braun (PSB 0582) |
| PMM191 | h-, leu1-32 ade6-210 ura4-D18 imr1L(Ncol)::ura4+ otr1R(SphI)::ade6+ mat1_m-cyhS smt0 rpl42::cyhR(sP56Q) cen1:hphMX, res2::KANMX | Fig. 1F; RT-QPCR | This study |
| PMM196 | h-, leu1-32 ade6-210 ura4-D18 imr1L(Ncol)::ura4+ otr1R(SphI)::ade6+ mat1_m-cyhS smt0 rpl42::cyhR(sP56Q) cen1:hphMX, res2::KANMX, pob3::NAT | Fig. 1F; RT-QPCR | This study |
| PMM41 | h-, leu1-32 ade6-210 ura4-D18 imr1L(Ncol)::ura4+ otr1R(SphI)::ade6+ mat1_m-cyhS smt0 rpl42::cyhR(sP56Q) cen1:hphMX, paf1::KANMX | Fig. 1E; RT-QPCR | This study |
| PMM47 | h-, leu1-32 ade6-210 ura4-D18 imr1L(Ncol)::ura4+ otr1R(SphI)::ade6+ mat1_m-cyhS smt0 rpl42::cyhR(sP56Q) cen1:hphMX, paf1::KANMX, pob3::NAT | Fig. 1E; RT-QPCR | This study |
| PMM133 | h-, leu1-32 ade6-210 ura4-D18 imr1L(Ncol)::ura4+ otr1R(SphI)::ade6+ mat1_m-cyhS smt0 rpl42::cyhR(sP56Q) cen1:hphMX, leo1::KANMX | Fig. 3B; RT-QPCR | This study |
| PMM135 | h-, leu1-32 ade6-210 ura4-D18 imr1L(Ncol)::ura4+ otr1R(SphI)::ade6+ mat1_m-cyhS smt0 rpl42::cyhR(sP56Q) cen1:hphMX, leo1::KANMX, pob3::NAT | Fig. 3B; RT-QPCR | This study |
| PMM72 | h-, smt0, ade6-M210, leu1-32, ura4-D18, locus2::mCherry::HygR, mat3M::ura4 | Fig. 3A, 3C-F, 2H, 4G; growth assays, RT-QPCR | Sigurd Braun (PSB0565) |
| PMM76 | h-, smt0, ade6-M210, leu1-32, ura4-D18, locus2::mCherry::HygR, mat3M::ura4, pob3::NAT | Fig. 3A, 3C-F, 2H, 4G; growth assays, RT-QPCR | This study |
| PMM370 | h-, smt0, ade6-M210, leu1-32, ura4-D18, locus2::mCherry::HygR, mat3M::ura4, leo1::KANMX | Fig. 3A; growth assays | This study |
| PMM375 | h-, smt0, ade6-M210, leu1-32, ura4-D18, locus2::mCherry::HygR, mat3M::ura4, leo1::KANMX, pob3::NAT | Fig. 3A; growth assays | This study |
| PMM357 | h-, smt0, ade6-M210, leu1-32, ura4-D18, locus2::mCherry::HygR, mat3M::ura4, epe1::KANMX | Fig. 3E, 3F, 3H; RT-QPCR, growth assays | This study |
| PMM360 | h-, smt0, ade6-M210, leu1-32, ura4-D18, locus2::mCherry::HygR, mat3M::ura4, epe1::KANMX, pob3::NAT | Fig. 3E, 3F, 3H; RT-QPCR, growth assays | This study |
| PMM363 | h-, smt0, ade6-M210, leu1-32, ura4-D18, locus2::mCherry::HygR, mat3M::ura4, mst2::KANMX | Fig. 3C, 3D; RT-QPCR, growth assays | This study |
| PMM366 | h-, smt0, ade6-M210, leu1-32, ura4-D18, locus2::mCherry::HygR, mat3M::ura4, mst2::KANMX, pob3::NAT | Fig. 3C, 3D; RT-QPCR, growth assays | This study |
| PMM385 | h90, cenH:: ade6p:SF-GFP (Kint2); mat3m(EcoRV)::ade6p:mKO2; ade6p:3xE2C:hygMX at Locus2; ΔREIII::REIII(Δs1, Δs2) | Fig. 2C, 2D, 2E, 4A-D, S2C-G; HSS assay, RT-QPCR, ChIP-QPCR, Western blot | Bassem Al-Sady (PAS332) |
| PAS331 | h90, cenH:: ade6p:SF-GFP (Kint2); mat3m(EcoRV)::ade6p:mKO2; ade6p:3xE2C:hygMX at Locus2; ΔREIII::REIII(Δs1, Δs2), clr4::KANMX | Fig. 2C; HSS assay | Bassem Al-Sady |
| PAS885 | h90, cenH:: ade6p:SF-GFP (Kint2); mat3m(EcoRV):: ade6p:mKO2; ade6p:3xE2C:hygMX at Locus2; ΔREIII::REIII(Δs1, Δs2, pob3::KANMX | Fig. 2C; HSS assay | Bassem Al-Sady |
| PMM388 | h90, cenH:: ade6p:SF-GFP (Kint2); mat3m(EcoRV):: ade6p:mKO2; ade6p:3xE2C:hygMX at Locus2; ΔREIII::REIII(Δs1, Δs2), KANMX::spt16-1 | Fig. 2D, 2E, 4A-D, S2C-G; HSS assay, RT-QPCR, ChIP-QPCR, Western blot, RT-QPCR, ChIP-QPCR | Bassem Al-Sady |
| PMM411 | h90, cenH:: ade6p:SF-GFP (Kint2); mat3m(EcoRV):: ade6p:mKO2; ade6p:3xE2C:hygMX at Locus2; ΔREIII::REIII(Δs1, Δs2), KANMX::spt16-1, epe1::NAT | Fig. 3H; HSS assay | This study |
| PMM433 | h90, cenH:: ade6p:SF-GFP (Kint2); mat3m(EcoRV):: ade6p:mKO2; ade6p:3xE2C:hygMX at Locus2; ΔREIII::REIII(Δs1, Δs2), clr4F449Y | Fig. 4C; RT-QPCR | This study |
| PMM439 | h90, cenH:: ade6p:SF-GFP (Kint2); mat3m(EcoRV):: ade6p:mKO2; ade6p:3xE2C:hygMX at Locus2; ΔREIII::REIII(Δs1, Δs2), clr4F449Y, KANMX::spt16-1 | Fig. 4C; RT-QPCR | This study |
| PMM518 | h-, ars1::prad15 cre-EBD-LEU2, H3.2 lox-HA-HYG-lox-T7, cdc25+, leu1-32 | Fig. 4G, S4D, S4E; RITE assay, ChIP-QPCR | This study |
| PMM549 | h-, ars1::prad15 cre-EBD-LEU2, H3.2 lox-HA-HYG-lox-T7, cdc25+, leu1-32, pob3::NAT | Fig. 4G, S4D, S4E; RITE assay, chip-QPCR | This study |
| PMM557 | h-, ars1::prad15 cre-EBD-LEU2, H3.2 lox-HA-HYG-lox-T7, cdc25+, leu1-32, pob3::NAT, epe1::KANMX | Fig. 4G, S4D, S4E; RITE assay, ChIP-QPCR | This study |
| PMM245 | h-, ade6-M216 htb1-FLAG::Hyg clr4-HA:KANMX | Fig. S2A; ChIP-QPCR | This study |
| PMM275 | h-, htb1-Flag::Hyg, pob3::NAT, clr4-HA::KANMX | Fig. S2A; ChIP-QPCR | This study |
| PMM321 | h-, ade6-M210 leu1-32 ura4-D18 | Fig. 4G; RT-QPCR | Daeyoung Lee (CBL13) |
| PMM322 | h-, fft3Δ::hphMX6 ade6-M210 leu1-32 ura4-D18 | Fig. 4E; RT-QPCR | Daeyoung Lee (CBL3306) |
| PMM323 | h-, spt16-1-kan ura4-D18 | Fig. 4E; RT-QPCR | Daeyoung Lee (CBL2461) |
| PMM325 | h-, fft3Δ::hphMX6 spt16-1-kan ade6-M210 leu1-32 ura4-D18 | Fig. 4E; RT-QPCR | Daeyoung Lee (CBL4056) |
| PMM378 | h-, smt0, ade6-M210, leu1-32, ura4-D18, locus2::mCherry::HygR, mat3M::ura4, clr3::KANMX | Fig. 4E; RT-QPCR | This study |
| PMM381 | h-, smt0, ade6-M210, leu1-32, ura4-D18, locus2::mCherry::HygR, mat3M::ura4, clr3::KANMX, pob3::NAT | Fig. 4E; RT-QPCR | This study |
| PMM445 | h+, leu1-32, ade6-210, ura4-DS/E(ura4-D18?), imr1L(NcoI)::ura4+, otrR(SphI)::ade6+, epe1-CBP-FLAG:KANMX | Fig. S3A, S3B; Western blot, ChIP-QPCR | Sigurd Braun (PSB2192) |
| PMM468 | h+, leu1-32, ade6-210, ura4-DS/E(ura4-D18?), imr1L(NcoI)::ura4+, otrR(SphI)::ade6+, epe1-CBP-FLAG:KANMX, pob3::NAT | Fig. S3A, S3B; Western blot, ChIP-QPCR | This study |
| PMM450 | h90, cenH:: ade6p:SF-GFP (Kint2); mat3m(EcoRV):: ade6p:mKO2; ade6p:3xE2C:hygMX at Locus2; ΔREIII::REIII(Δs1, Δs2), epe1-CBP-Flag:NAT | Fig. S3B; ChIP-QPCR | This study |
| PMM453 | h90, cenH:: ade6p:SF-GFP (Kint2); mat3m(EcoRV):: ade6p:mKO2; ade6p:3xE2C:hygMX at Locus2; ΔREIII::REIII(Δs1, Δs2), KANMX::spt16-1, epe1-CBP-Flag:NAT | Fig. S3B; ChIP-QPCR | This study |
| PAS99 | h<sup>+</sup>, ura4-D18, leu1-32, ade6-M216, his7-366 | Fig. S2B; cell cycle | Bassem Al-Sady |
| PAS475 | ars1::prad15 cre-EBD-LEU2, H3.2-lox-HA-HYG-lox-T7, cdc25<sup>ts</sup> | Fig. S2B; cell cycle | Karl Ekwall (HU2549) |
| PAS791 | h<sup>90</sup>, spt16-1:KANMX, ura4-D18 | Fig. S2B; cell cycle | Bassem Al-Sady |
| PAS886 | h<sup>+</sup>, pob3::KANMX | Fig. S2B; cell cycle | Bassem Al-Sady |

**Table S2.**
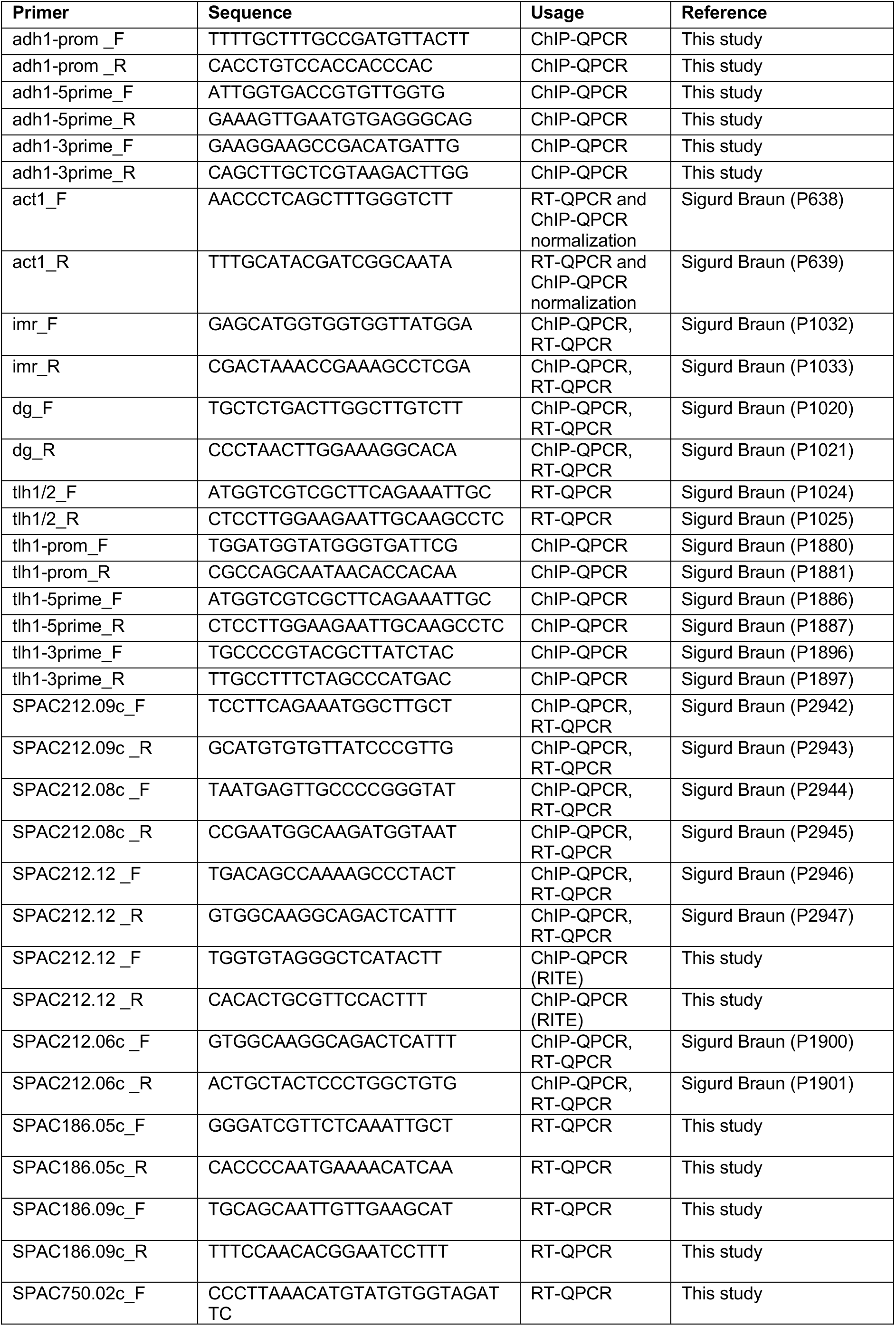
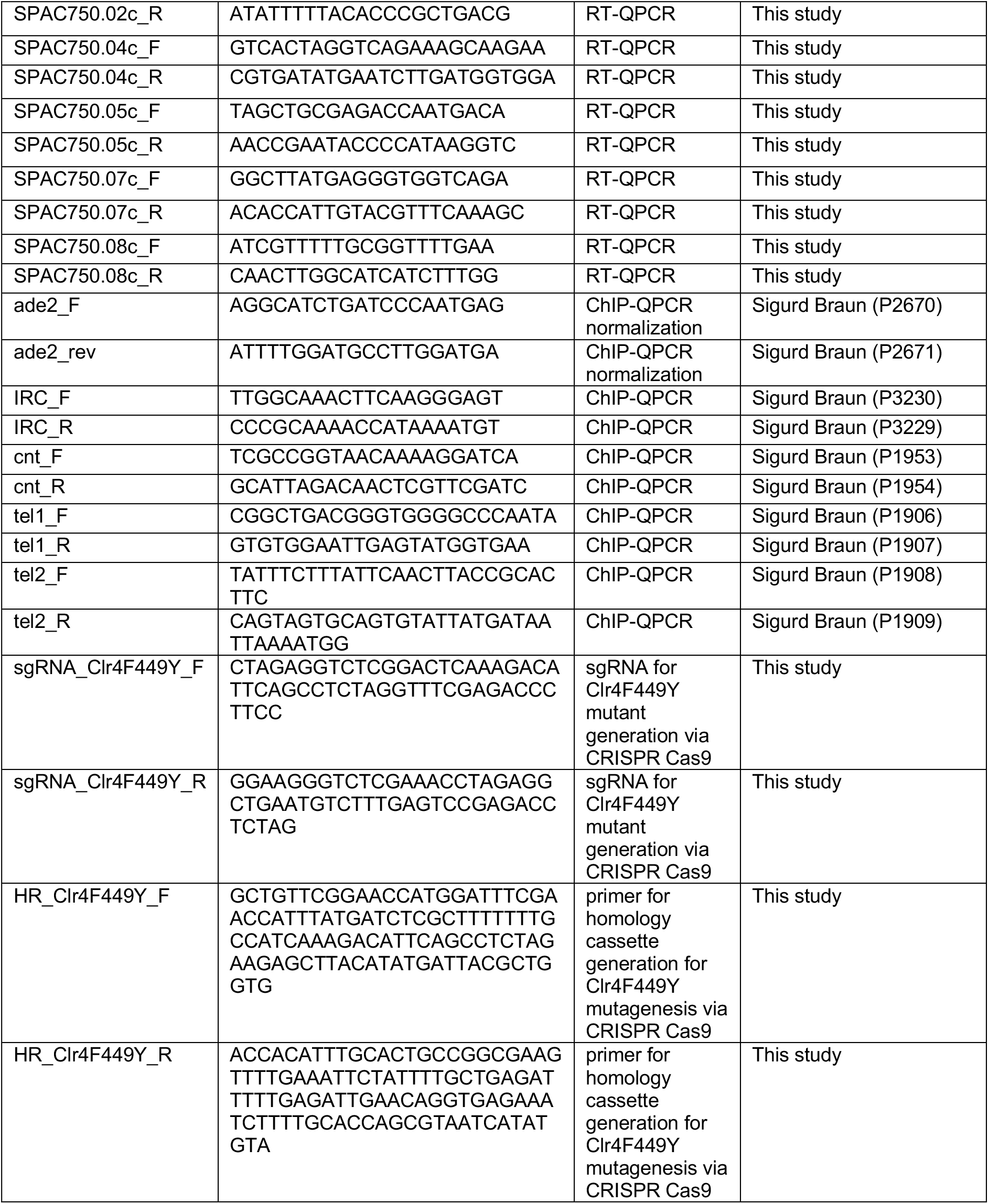
Primers used in this study

